# Do below-ground genotypes influence above-ground microbiomes of grafted tomato plants?

**DOI:** 10.1101/365023

**Authors:** Hirokazu Toju, Koji Okayasu, Michitaka Notaguchi

## Abstract

Bacteria and fungi form complex communities (microbiomes) in the phyllosphere and rhizosphere of plants, contributing to hosts’ growth and survival in various ways. Recent studies have suggested that host plant genotypes control, at least partly, microbial community compositions in the phyllosphere. However, we still have limited knowledge of how microbiome structures are determined in/on grafted crop plants, whose above-ground (scion) and below-ground (rootstock) genotypes are different with each other. By using eight varieties of grafted tomato plants, we examined how rootstock genotypes determine phyllosphere microbial assembly in field conditions. An Illumina sequencing analysis showed that both bacterial and fungal community structures did not significantly differ among tomato plants with different rootstock genotypes. Nonetheless, a further statistical analysis targeting respective microbial taxa suggested that some bacteria and fungi were preferentially associated with particular rootstock treatments. Specifically, a bacterium in the genus *Deinococcus* was found disproportionately from ungrafted tomato individuals. In addition, yeasts in the genus *Hannaella* were preferentially associated with the tomato individuals whose rootstock genotype was “Ganbarune”. Overall, this study suggests to what extent phyllosphere microbiome structures can be affected/unaffected by rootstock genotypes in grafted crop plants.

## INTRODUCTION

In both natural and agricultural ecosystems, bacteria and fungi in diverse taxonomic groups are associated with plants, positively and/or negatively influencing the survival and growth of their hosts (Vorholt 2012; Mendes et al. 2013; Bai et al. 2015; Peay et al. 2016). An increasing number of studies have shown that plant-associated microbes not only improve nutritional conditions of host plants but also increase plants’ resistance to abiotic stresses (e.g., high temperature, drought, and soil pollution) and that to pathogens and pests (Arnold et al. 2003; Mendes et al. 2011; Vandenkoornhuyse et al. 2015; Busby et al. 2017). In contrast, bacterial and fungal communities associated with plants can be regarded as serious risk factors in agriculture and forestry because they are occasionally dominated by plant pathogenic species or strains (Anderson et al. 2004; Callaway 2016). Therefore, controlling plant-associated microbiomes has been recognized as a major challenge towards the development of stable and sustainable management of crop fields and plantations (Schlaeppi & Bulgarelli 2015; Agler et al. 2016; Vorholt et al. 2017; Toju et al. 2018).

Host plant genotypes are among the most important factors determining microbiome structures (Whipps et al. 2008; Bodenhausen et al. 2014; Bulgarelli et al. 2015; Edwards et al. 2015). Developing disease-resistant crop plant varieties has been one of the major goals in breeding science (Collard & Mackill 2008; Dodds & Rathjen 2010; Dean et al. 2012). Moreover, recent studies have explored genes and mutations influencing whole microbiome structures (Hiruma et al. 2016; Castrillo et al. 2017), providing a basis for optimizing communities of plant-growth-promoting bacteria and/or fungi. Meanwhile, to gain more insights into mechanisms by which plant microbiomes are controlled, studies using plant individuals with complex genetic backgrounds have been awaited. Specifically, by using grafted plants, whose above- and below-ground genotypes are different with each other, we will be able to examine, for instance, how below-ground genetic factors control above-ground microbiome structures. Because root genotypes can control not only uptake of water and nutrients but also transport of phytohormones or signaling molecules (Goldschmidt 2014; Notaguchi & Okamoto 2015; Takahashi et al. 2018), their effects on leaf physiology potentially influence community compositions of endophytic and epiphytic microbes in the phyllosphere. Although studies focusing on such mechanisms interlinking above- and below-ground processes can provide essential insights into plants’ microbiome control, few attempts (Liu et al. 2018), to our knowledge, have been made to conduct experiments using grafted plants.

Grafting *per se* is a classic technique but it has been increasingly considered as a promising method for increasing yield, crop quality, abiotic stress resistance, and pathogen resistance of various plants (e.g., tomato, melon, grapevine, apple, and citrus) in agriculture (Khah et al. 2006; Martinez-Rodriguez et al. 2008; Flores et al. 2010; Rivard et al. 2012; Warschefsky et al. 2016). In general, performance of grafted plants depends greatly on compatibility between scion and rootstock genotypes (Ruiz & Romero 1999; Martinez-Ballesta et al. 2010; Schwarz et al. 2010). However, we still have limited knowledge of how scion–rootstock genotypic combinations determine microbiome structures in the phyllosphere and rhizosphere (Liu et al. 2018). Moreover, although some pioneering studies have investigated microbial community compositions of grafted plants (Ling et al. 2015; Song et al. 2015; Marasco et al. 2018), most of them focused on subsets of microbiomes (i.e., either bacteria or fungi but not both). Therefore, new lines of studies examining relationships between scion/rootstock genotypes and whole microbiome structures in roots/leaves have been awaited.

In this study, we evaluated how below-ground genotypes of plants determine bacterial and fungal community structures in/on leaves under field conditions. After growing grafted tomato [*Solanum lycopersicum* (= *Lycopersicon lycopersicum*)] individuals in a filed experiment, we analyzed the leaf microbial community compositions of the sampled tomatoes based on Illumina sequencing. The contributions of below-ground genotypes on the microbiome structures were then evaluated by comparing the microbial community datasets of eight tomato rootstock varieties. We also performed randomization-based statistical analyses to explore bacterial and fungal taxa that had strong signs of preferences for specific tomato rootstock varieties. Overall, this study suggests to what extent below-ground genotypes of plants influence above-ground plant–microbe interactions, providing a basis for managing microbiomes of grafted plants in agriculture and forestry.

## MATERIALS AND METHODS

### Grafted Tomato Seedlings

To prepare rootstocks, seeds of eight tomato varieties (“Chibikko”, “Ganbarune”, “M82”, “Micro-Tom”, “Regina”, “Spike”, “Triper”, and “Momotaro-Haruka”) were sown in 6-cm pots filled with potting soil on June 7, 2017 for “Momotaro-Haruka” and June 1, 2017 for the others, and then the pots were grown in a greenhouse of Togo Field, Nagoya University, Nagoya, Japan (35.112 ºN; 137.083 ºE). On June 22–23, seedlings for the field experiment detailed below were produced by grafting “Momotaro-Haruka” scions on each of the eight varieties of rootstocks: i.e., above-ground parts of the grafted seedlings were all Momotaro-Haruka, while below-ground parts differed among seedling individuals. Ungrafted “Momotaro-Haruka” seedlings were also prepared as control samples. The grafted (including Momotaro-Haruka/Momotaro-Haruka self-grafted seedlings) and ungrafted seedlings (in total, nine treatments) were grown in a greenhouse of Togo Field and, on July 7, they were transported to Center for Ecological Research, Kyoto University, Kyoto, Japan (34.972 ºN; 135.958 ºE). Each seedling was then transferred to a 9-cm pot filled with commercially-available culture soil (Rakuyo Co., Ltd.) on the day and they were kept on the field nursery shelf of Center for Ecological Research until the field experiment.

### Field Transplantation

On July 13, base fertilizer was provided to the soil in the experimental field of Center for Ecological Research (N = 13.6 g/m^2^; P_2_O_5_ = 13.6 g/m^2^; K_2_O = 13.6 g/m^2^). On July 25, the abovementioned seedlings (ca. 50 cm high) were transplanted to the open field at 50 cm horizontal intervals in three lines in a randomized order (9 seedling treatment × 5 replicates per line × 3 lines (sets) = 135 individuals; Fig. 1). The tomato individuals were watered twice (morning and evening) every day. On September 13, a ca. 1-cm^2^ disc of a mature leaf was sampled from each tomato individual and placed in a 2-mL microtube. The leaf samples were transferred to a laboratory of Center for Ecological Research using a cool box and they were then preserved at −80 ºC in a freezer until DNA extraction.

**FIGURE 1.**
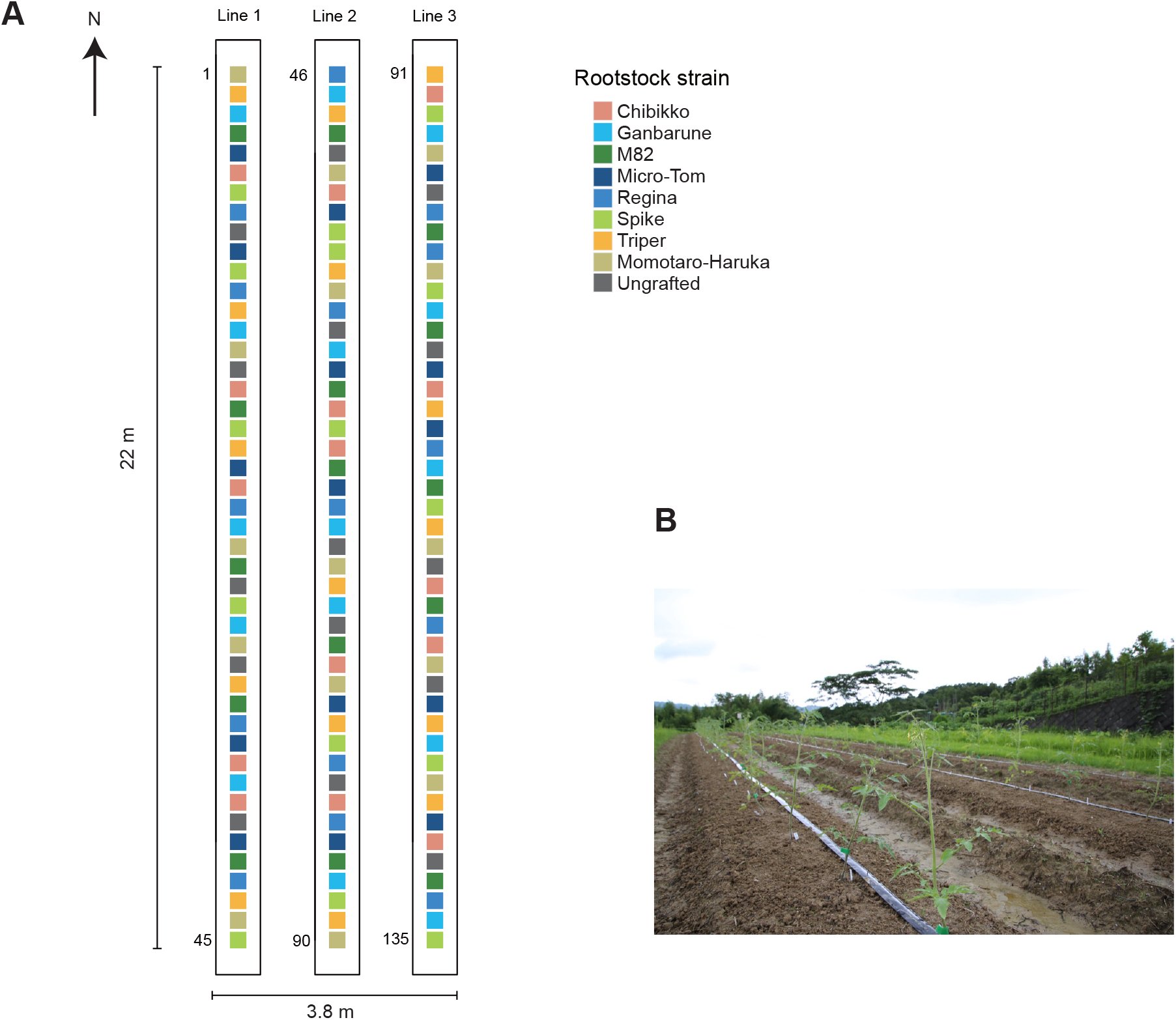
Field site. **(A)** Nine tomato rootstock varieties (treatments) in the field. For each rootstock variety, 15 replicate samples were transplanted to the field site (15 replicates × 9 varieties = 135 tomato individuals). The above-ground parts of all the 135 tomato individuals had the genotype of the tomato variety “Momotaro-Haruka”. **(B)** Transplanted tomato individuals.

### DNA Extraction, PCR, and Sequencing

Each leaf disc was surface-sterilized by immersing them in 1% NaClO for 1 min and it was subsequently washed in 70% ethanol. DNA extraction was extracted with a cetyltrimethylammonium bromide (CTAB) method after pulverizing the roots with 4 mm zirconium balls at 25 Hz for 3 min using a TissueLyser II (Qiagen).

For each leaf disc sample, the 16S rRNA V4 region of the prokaryotes and the internal transcribed spacer 1 (ITS1) region of fungi were PCR-amplified. The PCR of the 16S rRNA region was performed with the forward primer 515f (Caporaso et al. 2011) fused with 3– 6-mer Ns for improved Illumina sequencing quality (Lundberg et al. 2013) and the forward Illumina sequencing primer (5’- TCG TCG GCA GCG TCA GAT GTG TAT AAG AGA CAG- [3–6-mer Ns] – [515f] −3’) and the reverse primer 806rB (Apprill et al. 2015) fused with 3–6-mer Ns and the reverse sequencing primer (5’- GTC TCG TGG GCT CGG AGA TGT GTA TAA GAG ACA G [3–6-mer Ns] - [806rB] −3’) (0.2 μM each). To inhibit the PCR-amplification of mitochondrial and chloroplast 16S rRNA sequences of host plants, specific peptide nucleic acids [mPNA and pPNA; Lundberg et al. (2013)] (0.25 μM each) were added to the reaction mix of KOD FX Neo (Toyobo). The temperature profile of the PCR was 94 ºC for 2 min, followed by 35 cycles at 98 ºC for 10 s, 78 ºC for 10 s, 60 ºC for 30 s, 68 ºC for 50 s, and a final extension at 68 ºC for 5 min. To prevent generation of chimeric sequences, the ramp rate through the thermal cycles was set to 1 ºC/sec (Stevens et al. 2013). Illumina sequencing adaptors were then added to respective samples in the supplemental PCR using the forward fusion primers consisting of the P5 Illumina adaptor, 8-mer indexes for sample identification (Hamady et al. 2008) and a partial sequence of the sequencing primer (5’- AAT GAT ACG GCG ACC ACC GAG ATC TAC AC - [8-mer index] - TCG TCG GCA GCG TC −3’) and the reverse fusion primers consisting of the P7 adaptor, 8-mer indexes, and a partial sequence of the sequencing primer (5’- CAA GCA GAA GAC GGC ATA CGA GAT - [8-mer index] - GTC TCG TGG GCT CGG −3’). KOD FX Neo was used with a temperature profile of 94 ºC for 2 min, followed by 8 cycles at 98 ºC for 10 s, 55 ºC for 30 s, 68 ºC for 50 s (ramp rate = 1 ºC/s), and a final extension at 68 ºC for 5 min.

The PCR amplicons of the 135 tomato individuals (and negative control samples) were then pooled after a purification/equalization process with the AMPureXP Kit (Beckman Coulter). Primer dimers were removed from the pooled library by a supplemental AMpureXp purification process, in which the ratio of AMPureXP reagent to the pooled library was set to 0.6 (v/v).

The PCR of the fungal ITS1 region was performed with the forward primer ITS1F-KYO1 (Toju et al. 2012) fused with 3–6-mer Ns for improved Illumina sequencing quality (Lundberg et al. 2013) and the forward Illumina sequencing primer (5’- TCG TCG GCA GCG TCA GAT GTG TAT AAG AGA CAG- [3–6-mer Ns] – [ITS1F-KYO1] −3’) and the reverse primer ITS2-KYO2 (Toju et al. 2012) fused with 3–6-mer Ns and the reverse sequencing primer (5’- GTC TCG TGG GCT CGG AGA TGT GTA TAA GAG ACA G [3– 6-mer Ns] - [ITS2-KYO2] −3’). The PCR was performed based on the buffer and polymerase system of KOD FX Neo with a temperature profile of 94 ºC for 2 min, followed by 35 cycles at 98 ºC for 10 s, 58 ºC for 30 s, 68 ºC for 50 s, and a final extension at 68 ºC for 5 min. Illumina sequencing adaptors and 8-mer index sequences were added in the additional PCR and then the amplicons were purified and pooled as described above.

The sequencing libraries of the prokaryote 16S and fungal ITS regions were processed in an Illumina MiSeq sequencer (run center: KYOTO-HE; 15% PhiX spike-in). In general, quality of forward sequence data is generally higher than that of reverse sequence data in Illumina sequencing. Therefore, we optimized the settings of the Illumina sequencing run by targeting only forward sequences. Specifically, the numbers of the forward and reverse cycles were set 271 and 31, respectively: the reverse sequences were used only for discriminating between 16S and ITS1 sequences *in silico* based on the sequences of primer positions.

### Bioinformatics

The raw sequencing data were converted into FASTQ files using the Illumina’s program bcl2fastq 1.8.4. The obtained FASTQ files were demultiplexed with the program Claident v0.2.2018.05.29 (Tanabe & Toju 2013; Tanabe 2018), by which sequencing reads whose 8-mer index positions included nucleotides with low (< 30) quality scores were removed. The sequencing data were deposited to DNA Data Bank of Japan (DDBJ) (BioProject accession: PRJDB7150). Only forward sequences were used in the following analyses after trimming low-quality 3’-end sequences using Claident. Noisy reads (Tanabe 2018) were subsequently discarded and then denoised dataset consisting of 1,201,840 16S and 1,730,457 ITS1 reads were obtained.

For each region (16S or ITS1), filtered reads were clustered with a cut-off sequencing similarity of 97% using the program VSEARCH (Rognes et al. 2014) as implemented in Claident. The operational taxonomic units (OTUs) representing less than 10 sequencing reads were discarded and then the molecular identification of the remaining OTUs was performed based on the combination of the query-centric auto-*k*-nearest neighbor (QCauto) algorithm of reference database search (Tanabe & Toju 2013) and the lowest common ancestor (LCA) algorithm of taxonomic assignment (Huson et al. 2007) as implemented in Claident. Note that taxonomic identification results based on the QCauto-LCA pipeline are comparable to, or sometimes more accurate than, those with the alternative approaches (Tanabe & Toju 2013; Toju et al. 2016a; Toju et al. 2016b). In total, 143 prokatyote (bacterial or archaeal) OTUs and 529 fungal OTUs were obtained for the 16S and ITS1 regions, respectively (Supplementary Data 1). The UNIX codes used in the above bioinformatic pipeline are provided as Supplementary Data 2.

For each target region (16S or ITS1), we obtained a sample × OTU matrix, in which a cell entry depicted the number of sequencing reads of an OTU in a sample (Supplementary Data 3). To minimize effects of PCR/sequencing errors, cell entries whose read counts represented less than 0.1% of the total read count of each sample were removed [cf. Peay et al. (2015)]. The filtered matrix was then rarefied to 500 reads per sample using the “rrarefy” function of the vegan 2.4-5 package (Oksanen et al. 2017) of R 3.4.3 (R-Core-Team 2017). Samples with less than 500 reads were discarded in this process. In total, the rarefied matrices of the 16S and ITS1 regions included 125 and 132 samples, respectively: at least 13 replicate samples per treatment were retained in both datasets (Supplementary Data 4).

### Community Structure in the phyllosphere

Relationship between the number of sequencing reads and that of prokaryote/fungal OTUs was examined for each dataset (16S or ITS1) with the vegan “rarecurve” function of R. Likewise, relationship between the number of samples and that of OTUs was examined with the vegan “specaccum” function. For each dataset, difference in order- or genus-level community compositions among seedling treatments (rootstock varieties) was examined by the permutational analysis of variance [PERMANOVA; Anderson (2001)] with the vegan “adonis” function (10,000 permutations). To control spatial effects in the field experiment data, the information of replicate sample sets (Fig. 1) was included as an explanatory variable in the PERMANOVA. The “Raup-Crick” metric (Chase et al. 2011) was used to calculate *β*-diversity based on the order- or genus-level data matrices (Supplementary Data 5).

To explore prokaryote/fungal taxa whose occurrences on tomato individuals were associated with rootstock varieties, a series of analysis of variance (ANOVA) was performed. Specifically, based on the genus-level matrix of the 16S or ITS1 dataset (Supplementary Data 5), an ANOVA model was constructed for each prokaryote/fungal genus by including the proportion of the sequencing reads of the target genus and the rootstock variety information of host tomatoes as response and explanatory variables, respectively. The information of replicate samples (i.e., location information) was included as an additional explanatory variable. Genera that occurred in less than 30 tomato individuals were excluded from the analysis.

### Randomization Analysis of Preferences for Rootstock Varieties

We further explored prokaryote/fungal taxa showing preferences for specific rootstock varieties based on a randomization analysis. In the sample × genus matrix of the 16S or ITS1 dataset (Supplementary Data 5), the labels of rootstock varieties were shuffled (100,000 permutations) and then preference of a prokaryote/fungal genus (*i*) for a rootstock variety (*j*) was evaluated as follows:

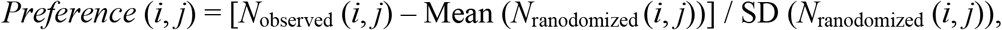

where *N*_observed_ (*i*, *j*) denoted the mean number of the sequencing reads of genus *i* across rootstock variety *j* tomato samples in the original data, and the Mean (*N*_ranodomized_ (*i*, *j*)) and SD (*N*_ranodomized_ (*i*, *j*)) were the mean and standard deviation of the number of sequencing reads for the focal genus–rootstock combination across randomized matrices. Genera that occurred in 30 or more tomato individuals were subjected to the randomization analysis.

For the genera that showed significant preferences for specific tomato rootstock varieties, we performed an additional analysis to evaluate which bacterial/fungal OTUs in each genus had strong host-variety preferences. Specifically, the randomization analysis of the above preference index (100,000 permutations) was applied to rarefied sample × OTU matrix of the 16S or ITS1 dataset (Supplementary Data 4). OTUs that occurred in less than 30 tomato individuals were excluded from the analysis.

## RESULTS

### Community Structure in the phyllosphere

On average, 13.6 (SD = 4.2) prokaryote and 26.3 (SD = 9.4) fungal OTUs per sample were observed in the rarefied data matrices (Supplementary Fig. 1). The total numbers of prokaryote and fungal OTUs included in the rarefied datasets were 116 and 413, respectively (Supplementary Data. 4). All the prokaryote OTUs belonged to Bacteria: no archaeal OTUs were observed.

In the bacterial community of the tomato phyllosphere, bacteria in the orders Sphingomonadales and Rhizobiales were dominant (Fig. 2A). Bacteria in the order Pseudomonadales were frequently observed, too, across the tomato varieties examined. Meanwhile, bacteria in the order Deinococcales were abundant only in the ungrafted tomato individuals (Fig. 2A). At the genus-level, the genera *Sphingomonas*, *Methylobacterium*, and *Pseudomonas* were frequently observed across the rootstock varieties examined, while *Deinococcus* bacteria were abundant only in the ungrafted tomatoes (Fig. 2B).

**FIGURE 2.**
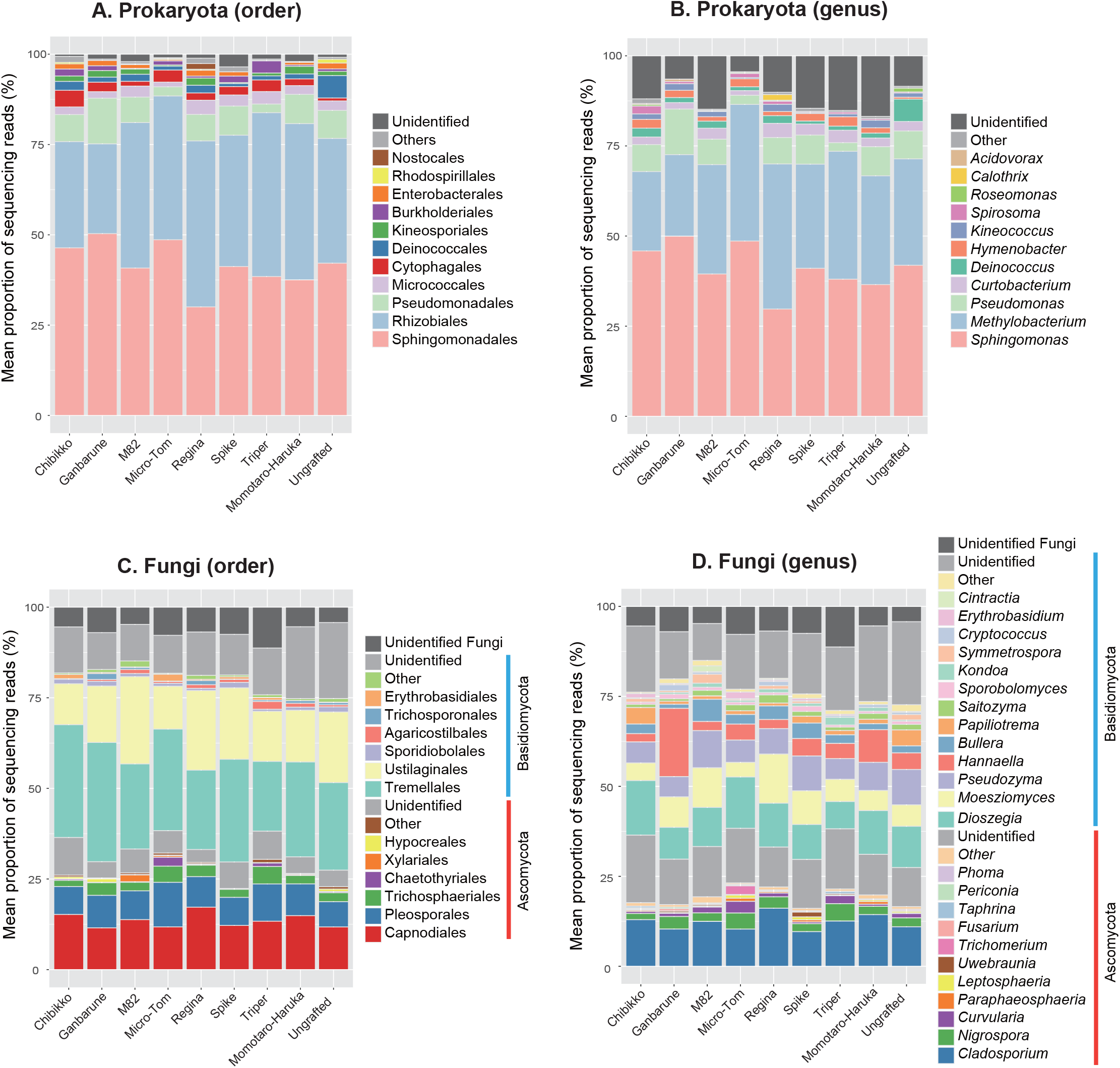
Structure of the phyllosphere microbial communities. The phyllosphere microbial community compositions were compared among tomato individuals with different rootstock genotypes. **(A)** Order-level community structure of prokaryotes. **(B)** Genus-level community structure of prokaryotes. **(C)** Order-level community structure of fungi. **(D)** Genus-level community structure of fungi.

In the phyllosphere fungal community, ascomycete fungi in the orders Capnodiales and Plesporales and the basidiomycete fungi in the orders Tremellales and Ustiaginales were abundant (Fig. 2C). At the genus-level, *Cladosporium*, *Dioszegia*, *Moesziomyces* (anamorph = *Pseudozyma*), and *Hannaella* were frequently observed (Fig. 2D). Among them, *Hannaella* fungi dominated the phyllosphere fungal community of the tomato rootstock variety “Ganbarune” (the proportion of *Hannaella* reads = 19.0 %), while their proportion was relatively low on other host varieties (2.3–9.1 %; Fig. 2D).

A statistical test based on PERMANOVA showed that replicate sampling positions, but not tomato rootstock varieties, significantly explained variation in the whole structure of the bacterial/fungal community (Table 1). However, further analyses targeting respective genera (Table 2 and 3) indicated that the proportion of the fungal genus *Hannaella* varied among tomato rootstock varieties, although the pattern was non-significant after a Bonferroni correction of *P* values. Meanwhile, the proportion of some taxa such as the bacterial genus *Sphingomonas* and the fungal genus *Cladosporium* varied significantly among replicates (Tables 2 and 3), suggesting that spatial positions in the experimental field affected the formation of the phyllosphere microbial communities of the tomato plants.

**TABLE 1.** Effects of rootstock varieties and spatial positions on the entire microbial community structure. A PERMANOVA was conducted for each target community (prokaryotes or fungi) at each taxonomic level (order or genus). The rootstock varieties of host tomato and spatial positions in the field (location; Fig. 1A) were considered as explanatory variables.

| Taxon | Taxonomic level | Variable | df | $F_{\text{model}}$ | $R^2$ | $P$ |
| --- | --- | --- | --- | --- | --- | --- |
| Prokaryotes | Order | Variety | 8 | 1.0 | 0.061 | 0.4731 |
|  |  | Location | 14 | 1.6 | 0.173 | 0.0379 |
|  | Genus | Variety | 8 | 1.1 | 0.064 | 0.3733 |
|  |  | Location | 14 | 2.1 | 0.207 | 0.0035 |
| Fungi | Order | Variety | 8 | 0.6 | 0.033 | 0.7509 |
|  |  | Location | 14 | 2.2 | 0.213 | 0.0119 |
|  | Genus | Variety | 8 | 0.9 | 0.050 | 0.5586 |
|  |  | Location | 14 | 1.9 | 0.185 | 0.0350 |

**TABLE 2.** Effects of rootstock varieties and spatial positions on the proportion of each prokaryote genus in the community data. For each prokaryote genus, an ANOVA model of the mean proportion of sequencing reads was constructed by including the rootstock varieties of host tomato and spatial positions in the field (location; Fig. 1A) as explanatory variables. Genera that occurred in 30 or more tomato individuals were subjected to the analysis.

| Genus | Variety |  |  | Location |  |  |
| --- | --- | --- | --- | --- | --- | --- |
|  | df | *F* | *P* | df | *F* | *P* |
| *Curtobacterium* | 8 | 0.3 | 0.9710 | 14 | 1.1 | 0.3260 |
| *Deinococcus* | 8 | 1.8 | 0.0944 | 14 | 1.3 | 0.2386 |
| *Hymenobacter* | 8 | 0.5 | 0.8730 | 14 | 1.1 | 0.3900 |
| *Kineococcus* | 8 | 0.7 | 0.6710 | 14 | 0.6 | 0.8970 |
| *Methylobacterium* | 8 | 1.7 | 0.0986 | 14 | 2.0 | 0.0229 |
| *Pseudomonas* | 8 | 1.7 | 0.1060 | 14 | 0.6 | 0.8490 |
| *Sphingomonas* | 8 | 2.0 | 0.0538 | 14 | 3.2 | 0.0004 |
| *Spirosoma* | 8 | 1.0 | 0.4230 | 14 | 1.0 | 0.4310 |

**TABLE 3.**
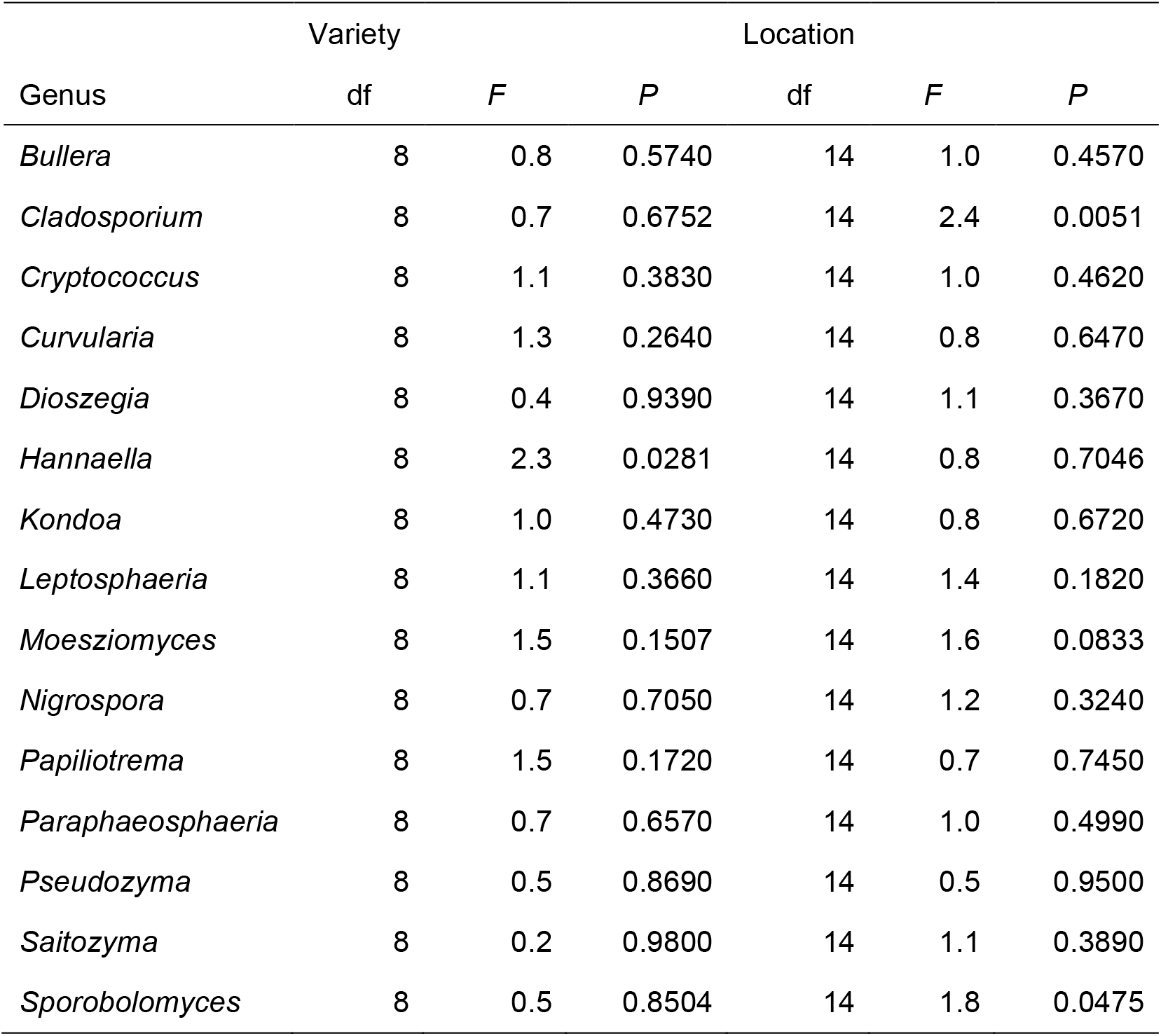
Effects of rootstock varieties and spatial positions on the proportion of each fungal genus in the community data. For each fungal genus, an ANOVA model of the mean proportion of sequencing reads was constructed by including the rootstock varieties of host tomato and spatial positions in the field (location; Fig. 1A) as explanatory variables. Genera that occurred in 30 or more tomato individuals were subjected to the analysis.

| Genus | Variety |  |  | Location |  |  |
| --- | --- | --- | --- | --- | --- | --- |
|  | df | *F* | *P* | df | *F* | *P* |
| *Bullera* | 8 | 0.8 | 0.5740 | 14 | 1.0 | 0.4570 |
| *Cladosporium* | 8 | 0.7 | 0.6752 | 14 | 2.4 | 0.0051 |
| *Cryptococcus* | 8 | 1.1 | 0.3830 | 14 | 1.0 | 0.4620 |
| *Curvularia* | 8 | 1.3 | 0.2640 | 14 | 0.8 | 0.6470 |
| *Dioszegia* | 8 | 0.4 | 0.9390 | 14 | 1.1 | 0.3670 |
| *Hannaella* | 8 | 2.3 | 0.0281 | 14 | 0.8 | 0.7046 |
| *Kondoa* | 8 | 1.0 | 0.4730 | 14 | 0.8 | 0.6720 |
| *Leptosphaeria* | 8 | 1.1 | 0.3660 | 14 | 1.4 | 0.1820 |
| *Moesziomyces* | 8 | 1.5 | 0.1507 | 14 | 1.6 | 0.0833 |
| *Nigrospora* | 8 | 0.7 | 0.7050 | 14 | 1.2 | 0.3240 |
| *Papiliotrema* | 8 | 1.5 | 0.1720 | 14 | 0.7 | 0.7450 |
| *Paraphaeosphaeria* | 8 | 0.7 | 0.6570 | 14 | 1.0 | 0.4990 |
| *Pseudozyma* | 8 | 0.5 | 0.8690 | 14 | 0.5 | 0.9500 |
| *Saitozyma* | 8 | 0.2 | 0.9800 | 14 | 1.1 | 0.3890 |
| *Sporobolomyces* | 8 | 0.5 | 0.8504 | 14 | 1.8 | 0.0475 |

### Randomization Analysis of Preferences for Rootstock Varieties

A randomization analysis indicated that the bacterial genus *Deinococcus* occurred preferentially on the ungrafted tomato individuals (Fig. 3A). Likewise, the fungal genus *Hannaella* showed preferences for the rootstock variety “Ganbarune” (Fig. 3B). In an additional randomization analysis, a bacterial OTU phylogenetically allied to *Deinococcus citri* (P_040) and fungal OTUs allied to *Hannaella oryzae* (F_427 and F_428) displayed statistically significant preferences for ungrafted and “Ganbarune” tomato plants, respectively (Table 4).

**FIGURE 3.**
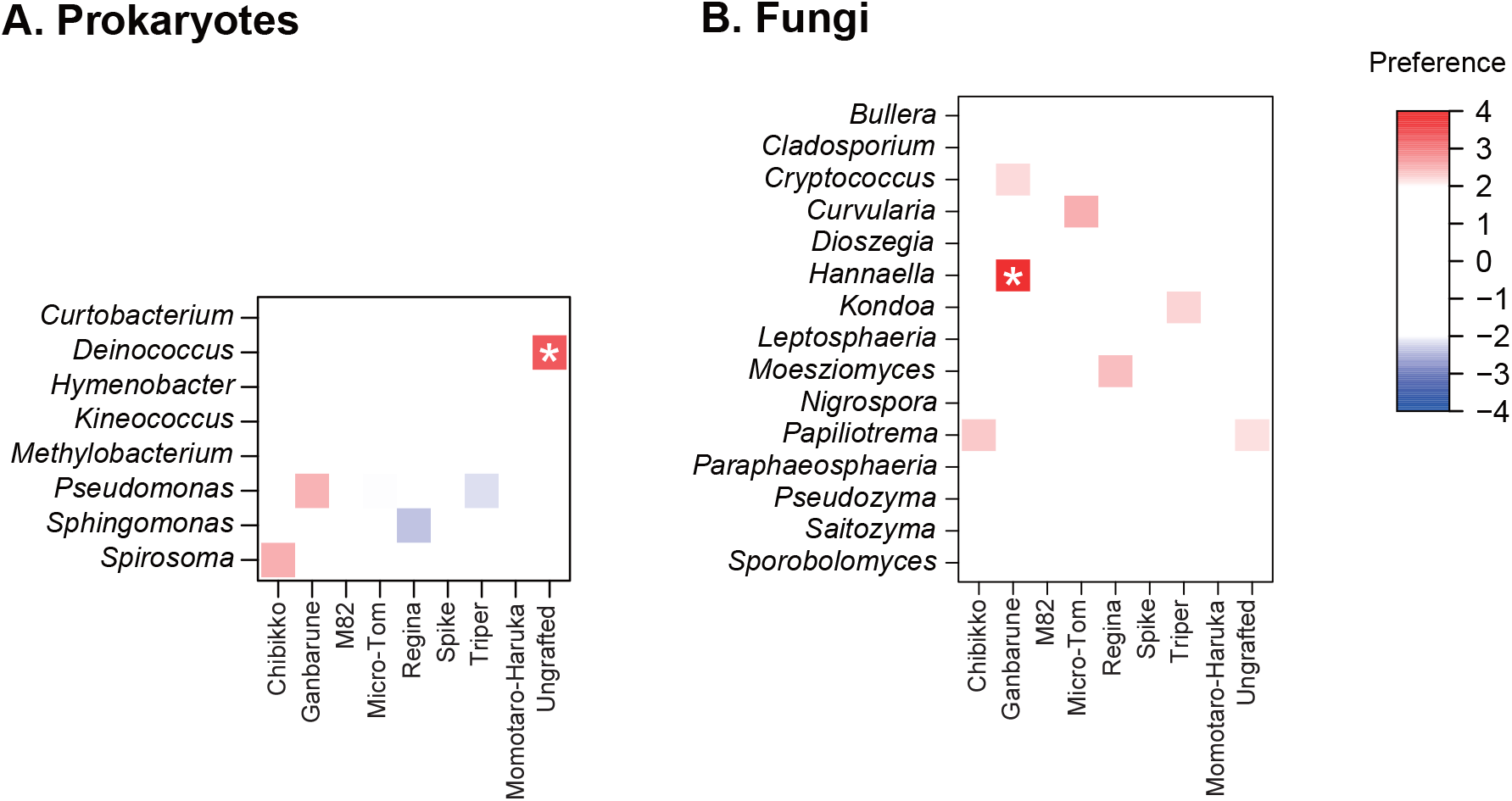
Randomization analysis of preferences for rootstock varieties. An asterisk indicates significant preference index score in a combination of a microbial genus and a host rootstock variety (Bonferroni correction applied to each genus; *α* = 0.05). **(A)** Prokatyote genera. **(B)** Fungal genera.

**TABLE 4.** Prokaryote and fungal OTUs showing statistically significant preferences for tomato rootstock varieties.

| OTU | Preferred variety | Phylum | Class | Order | Family | Genus | NCBI Blast top hit | Accession | Cover | Identity |
| --- | --- | --- | --- | --- | --- | --- | --- | --- | --- | --- |
| Prokaryotes |  |  |  |  |  |  |  |  |  |  |
| P_040 | Ungrafted ($P = 0.00321$) | Deinococcus-Thermus | Deinococci | Deinococcales | Deinococcaceae | *Deinococcus* | *Deinococcus citri* | LT602922 | 100% | 100% |
| Fungi |  |  |  |  |  |  |  |  |  |  |
| F_427 | Ganbarune ($P = 0.00078$) | Basidiomycota | Tremellomycetes | Tremellales | Bulleribasidiaceae | *Hannaella* | *Hannaella oryzae* | KY103504 | 89% | 99% |
| F_428 | Ganbarune ($P = 0.00099$) | Basidiomycota | Tremellomycetes | Tremellales | Bulleribasidiaceae | *Hannaella* | *Hannaella oryzae* | KY103504 | 89% | 99% |

## DISCUSSION

The field experiment using eight tomato rootstock varieties suggested that below-ground plant genotypes did not significantly affect the entire structures of the phyllosphere microbiomes (Table 1). However, detailed analyses indicated the existence of phyllosphere microbial taxa whose associations with host plants were affected by below-ground plant genotypes (Figs. 2 and 3; Tables 2–4). Thus, this study not only shows to what extent above-ground microbiome structures of grafted plants are affected/unaffected by below-ground genotypes but also suggests which phyllosphere microbial taxa can be managed by selecting rootstock varieties of crop plants.

The phyllosphere bacterial communities of the tomato individuals analyzed in this study were dominated by Alphaproteobacteria (e.g., *Sphingomonas* and *Methylobacterium*) as well as Gammaproteobacteria (e.g., *Pseudomonas*) as has been reported in previous studies on crop and non-crop plants (Lindow & Brandl 2003; Vorholt 2012; Bai et al. 2015) (Fig. 2). Among the dominant bacteria, *Pseudomonas* is recognized mainly as plant pathogenic taxon (Buell et al. 2003; Yu et al. 2013), although some *Pseudomonas* species are known to suppress leaf fungal pathogens by producing antibiotics (Flaishman et al. 1996; De Meyer & Höfte 1997). The genus *Sphingomonas* is known to involve species that protect host plants against *Pseudomonas* pathogens (Innerebner et al. 2011; Vogel et al. 2012) or promote plant growth by producing phytohormones such as gibberellins and indole acetic acid (Khan et al. 2014). Bacteria in the genus *Methylobacterium* are often localized around stomatal pores in the phyllosphere (Abanda-Nkpwatt et al. 2006), using plant-derived methanol as principal carbon source (Delmotte et al. 2009; Schauer & Kutschera 2011; Knief et al. 2012; Ryffel et al. 2016). Genomic studies have shown that *Methylobacterium* genomes involve genes of metabolic pathways that potentially contribute to host plant growth (e.g., auxin biosysnthesis, cytokine biosynthesis, and vitamin B_12_ biosynthesis) (Kwak et al. 2014). *Methylobacterium* is also known to induce resistance of plants against fungal pathogens, nominated as prospective a biocontrol agent (Madhaiyan et al. 2006). Thus, these dominant bacteria, whose associations with hosts are likely irrespective of host below-ground genotypes (Fig. 2), may affect growth of tomato plants both positively and negatively.

Our data also indicated that fungi in the ascomycete genus *Cladosporium* and the basidiomycete genera *Dioszegia* and *Moesziomyces* (anamorph = *Pseudozyma*) were abundant within the tomato phyllosphere (Fig. 2). Among them, *Cladosporium* involves a well-characterized pathogenic species, *C. fulvum*, which causes tomato leaf mold (De Wit & Spikman 1982; van Kan et al. 1991; Jones et al. 1994; Rivas & Thomas 2005). The basidiomycete taxa listed above are characterized by their anamorphic yeast forms and they have been observed in the phyllosphere of various plant species (Inácio et al. 2005; Karlsson et al. 2014; Sapkota et al. 2015; Kruse et al. 2017). For example, *Dioszegia*, a basidiomycete taxon in the order Tremellales, has been reported from cereal and *Arabidopsis* (Sapkota et al. 2015; Wang et al. 2016), potentially playing key roles within microbe–microbe interaction webs in the phyllosphere (Agler et al. 2016). The genus *Moesziomyces* is represented by plant-pathogenic smut fungi (Diagne-Leye et al. 2013). However, a recent phylogenetic study of teleomorphic (*Moesziomyces*) and anamorphic (*Pseudozyma*) specimens (Kruse et al. 2017) suggested that this Ustilaginaceae taxon could involve not only phytopathogenic species but also species with antifungal properties against the causal agent of cucumber powdery mildew (*Podosphaera fuliginea*) (Avis et al. 2001) or those that can induce resistance of host plants against fungal pathogens such as *Botrytis cinerea* (Buxdorf et al. 2013). Thus, the community data, as a whole, suggest that not only dominant bacterial taxa but also various fungal taxa potentially play complex physiological roles in the phyllosphere of tomato plants.

While there were bacterial and fungal taxa commonly associated with tomato plants irrespective of host below-ground genotypes, fungi in the genus *Hannaella* displayed preferences for rootstock genotypes (Fig. 3; Tables 3 and 4). Specifically, *Hannaella* was the most abundant fungal taxon in the tomato individuals whose rootstock genotype was “Ganbarune” (Fig. 2). Like other yeast taxa in Tremellaceae (e.g., *Derxomyces* and *Dioszegia*) (Wang & Bai 2008), *Hannaella* yeasts are frequently observed in the phyllosphere of various plant species (Nutaratat et al. 2014; Kaewwichian et al. 2015; Nasanit et al. 2015; Nasanit et al. 2016). Some *Hannaella* species are known to produce indol acetic acid (Nutaratat et al. 2014; Sun et al. 2014), although a study has suggested that the yeasts do not necessarily promote plant growth (Sun et al. 2014). Therefore, it remains a challenge to understand how *Hannaella* yeasts interact with other yeasts and bacterial/fungal species in/on plant leaves and how they influence plant performance host-genotype specifically.

The randomization analysis performed in this study also indicated that a bacterial OTU phylogenetically allied to the *Deinococcus* species isolated from leaf canker lesions of citrus trees (Ahmed et al. 2014) had a preference for ungrafted tomato individuals (Fig. 3; Tables 2 and 4). Given that this bacterial OTU was rarely observed in self-grafted tomato individuals (Fig. 2), grafting treatment *per se*, rather than plant genotypes, could be responsible for the biased distribution of the bacterium. This finding is of particular interest because *Deinococcus* is famous for its high tolerance to desiccation (Mattimore & Battista 1996; Tanaka et al. 2004). Grafting itself has been recognized as a way for increasing plants’ resistance against drought stress (Schwarz et al. 2010; Warschefsky et al. 2016). Thus, the above-ground parts of the ungrafted tomato plants might uptake less water than grafted plants, resulting in the high proportion of the desiccation-tolerant bacteria in the phyllosphere.

Although this study provides some implications for how phyllosphere microbiomes of grafted plants can be influenced by rootstock genotypes, potential pitfalls of the present results should be taken into account. First, as our data were based on a snapshot sampling in the late growing season of tomato, we are unable to infer the timing at which the observed bacteria and fungi colonized the tomato phyllosphere. Therefore, some of the detected bacterial and fungal OTUs might colonize the tomato individuals before they were transplanted into the experimental field. However, given that spatial positions within the field had significant effects on the microbial community structures (Table 1), colonization of indigenous (resident) microbes in the field could be a major factor determining the observed microbiome pattern. Second, we need to acknowledge that microbiome profiling with high-throughput DNA sequencing *per se* does not reveal the fine-scale distribution of the detected microbial OTUs in the phyllosphere. Although we surface-sterilized the leaf samples, the microbiome data involved not only possibly endophytic taxa but also bacteria and fungi that have been regarded as epiphytes (e.g., *Methylobacterium*) (Omer et al. 2004; Abanda-Nkpwatt et al. 2006) [but see Jourand et al. (2004)]. Microscopic analyses with taxon-specific fluorescent probes, for example, will provide essential insights into the localization of the observed microbes in the phyllosphere. Third, while this study was designed to examine effects of below-ground genotypes on above-ground parts of grafted plants, recent studies have shown that genetic materials (i.e., DNA) can be transported between scion and rootstock tissue, at least at graft junction region, in a grafted plant (Stegemann & Bock 2009). Thus, contributions of above-/below-ground genotypes to rhizosphere/phyllosphere microbiomes may be much more complex than had been assumed in this study.

Overall, this study suggested that majority of phyllosphere microbes can be associated with grafted tomato plants irrespective of rootstock genotypes of their hosts. Meanwhile, phyllosphere microbial taxa could display preferences for grafted/ungrafted plants or specific host rootstock varieties. Both grafting and the use of plant-beneficial microbes have been regarded as prospective options for securing agricultural/forestry production in the era of increasing biotic and abiotic environmental stresses (Schwarz et al. 2010; Schlaeppi & Bulgarelli 2015; Warschefsky et al. 2016; Toju et al. 2018). Further integrative studies will help us explore best conditions in which grafting and microbiome technologies are merged into a solid basis of stable and sustainable agricultural practices.

## AUTHOR CONTRIBUTIONS

H.T., K.O., and M.N. performed experiments. H.T. analyzed data. H.T. wrote the paper with M.N.

## ACKNOWLEDGEMENTS

We thank Satomi Yoshinami, Akira Matsumoto, Tomoaki Muranaka, Sarasa Amma, and Hiroki Kawai for their support in field experiment and/or molecular experiments. This work was financially supported by JST PRESTO (JPMJPR16Q6) to H.T. and MAFF science and technology research promotion program for agriculture, forestry, fisheries and food industry grant (16770567) and JST PRESTO (JPMJPR15O3) to M.N..

## SUPPLEMENTARY MATERIAL

The Supplementary Material for this article can be found online at: XXXX.

